# BrainAGE: Revisited and reframed machine learning workflow

**DOI:** 10.1101/2022.11.21.517386

**Authors:** Polona Kalc, Robert Dahnke, Felix Hoffstaedter, Christian Gaser, Alzheimer’s Disease Neuroimaging Initiative

## Abstract

Since the introduction of the BrainAGE method (Franke et al., 2010), novel machine learning methods of brain age prediction have continued to emerge. The idea of estimating the chronological age from magnetic resonance images proved to be an interesting field of research due to the relative simplicity of its interpretation and its potential use as a biomarker of brain health. We revised our previous BrainAGE approach, originally utilising relevance vector regression (RVR), and substituted it with Gaussian process regression (GPR), which enables more stable processing of larger datasets, such as the UK Biobank (UKB). In addition, we extended the global BrainAGE approach to regional BrainAGE, providing spatially specific scores for 5 brain lobes per hemisphere. We tested the performance of the new algorithms under several different conditions and investigated their validity on the ADNI and schizophrenia samples, as well as on a synthetic dataset of neocortical thinning. The results show an improved performance of the reframed global model on the UKB sample with a mean absolute error (MAE) of less than 2 years and a significant difference in BrainAGE between healthy participants and patients with Alzheimer’s disease and schizophrenia. Moreover, the workings of the algorithm show meaningful effects for a simulated neocortical atrophy dataset. The regional BrainAGE model performed well on two clinical samples, showing disease-specific patterns for different levels of impairment. The results demonstrate that the new improved algorithms provide reliable and valid brain age estimations.

## Introduction

Our original BrainAGE framework (Franke et al., 2010) is a supervised machine learning method that employs relevance vector regression (RVR) in order to predict the chronological age of an individual based on a single T1-weighted magnetic resonance (MR) scan of their brain. The difference between the apparent age of the individual’s brain as predicted by the algorithm and their chronological age is the so-called brain age gap estimation (BrainAGE). A positive gap implies more prominent structural brain changes that commonly occur with ageing progression, e.g. decrease of grey matter volume (Bethlehem et al., 2022), whereas a negative difference signifies a slower ageing in comparison to other subjects within a sample. The BrainAGE method was introduced more than a decade ago and was successfully applied to various subject groups, such as adolescents (Franke et al., 2012), elderly people with mild cognitive impairment (MCI) and Alzheimer’s disease (AD; Franke et al., 2010; Franke & Gaser, 2012), musicians (Rogenmoser et al., 2018), meditators (Luders et al., 2016), and even other animal species, such as rodents and non-human primates (Franke et al., 2017). The method proved to be a reliable biomarker of AD (Franke & Gaser, 2012), Parkinson’s disease (Eickhoff et al., 2021), and is associated with brain alterations in schizophrenia (Nenadić et al., 2017) and type 2 diabetes (Franke et al., 2013). A complete review can be found in Franke & Gaser (2019) or Cole & Franke (2017).

Since the introduction of the concept of brain age as a biomarker of ageing brains, numerous machine learning and deep learning models of estimating the age of the brain based on different MR imaging modalities have emerged (e.g., Beheshti et al., 2020; Cole et al., 2018; Hahn et al., 2022; Smith et al., 2019; Varzandian et al., 2021). The advancement of deep learning methods in the field of brain age prediction (e.g., Abrol et al., 2021; Gong et al., 2020; Hahn et al., 2022; Lombardi et al., 2021; Peng et al., 2021) and the availability of large multimodal imaging data collection, such as the UK Biobank (UKB; https://www.ukbiobank.ac.uk/), pushed the performance of the brain age prediction to unprecedented levels, managing to decrease mean absolute errors (MAE; i.e., a metric commonly used to compare the accuracy of prediction) to as low as 2.14 years (Peng et al., 2021).

The aim of the study was to revise the original BrainAGE machine learning framework to be able to keep pace with the advancement of deep learning algorithms in dealing with large datasets, while retaining the parsimonious and straightforward nature of voxel-based morphometry (VBM). Studies testing the performance of multiple algorithms in brain age estimation showed a good performance of Gaussian process regression (GPR) (Baecker et al., 2021; Beheshti et al., 2022; Cole et al., 2015; Cole et al., 2018; More et al. 2023), which can be seen as an alternative approach to artificial neural networks (Chen & Ren, 2009). We therefore implemented GPR in the new BrainAGE framework and utilised model ensembling to limit overfitting and improve prediction accuracy (van Veen et al., 2015). We replaced the initial BrainAGE framework (Franke et al., 2010; Franke & Gaser, 2012) with a combination of models using grey- or white matter tissue segments (of varying spatial resolutions and smoothing sizes) associated by (weighted) averaging and stacking. In addition to the global BrainAGE, we further implemented a regional BrainAGE approach to obtain greater spatial specificity for future applications to various clinical groups.

In order to evaluate the performance of our novel BrainAGE algorithms and to allow for straightforward comparison to results of other groups, we first focused on the UK Biobank (UKB) cohort. The UKB is a very large biomedical database and research resource that contains genetic, lifestyle and health information from half a million UK Biobank participants. We investigated the effects of various conditions (e.g., regularisation, varying training sample size) on prediction accuracy and examined the reliability of the predictions. In the second part of the paper, we tested the performance of the global and regional models on external datasets. We applied the new global model to a novel synthetic dataset of global neocortical thinning (Rusak et al., 2022), and to two samples from clinical populations, namely MCI and AD patients from the ADNI cohort as well as schizophrenia patients (SZ).

## Method

### Datasets

This research has been conducted using data from UK Biobank under the application number 41655. The UK Biobank has ethical approval from the North West Multi-Centre Research Ethics Committee (MREC) and is in possession of informed consents from the study cohort participants. Participants with pre-existing neurological or psychiatric diagnoses were excluded from the analyses (UKB data-fields #41202-0.0 to 41202-0.78 Diagnoses - main ICD10: presence of F0-F99 Mental and behavioural disorders). Overall, we analysed 36840 T1-weighted MR scans (52% women; age range: 45–82; *M*_Age_ = 64.09, *SD*_Age_ = 7.53 years). Some of the analyses were run on a pseudo random subsample of 7241 scans (52% women; age range: 45–82; *M*_Age_ = 64.08, *SD*_Age_ = 7.54 years), including only subject IDs starting with 1* to avoid the introduction of selection bias. For the cross-site prediction, we randomly sampled a fifth of the full sample (*n* = 7368) to obtain two size-matched subsamples from the scanning site 1 (Cheadle) and the scanning site 3 (Newcastle). The tests performed on the UKB sample served as a benchmark to compare our different BrainAGE models to the ones from other groups (e.g., Baecker et al., 2021; Peng et al., 2021).

Although the UKB is one of the largest neuroimaging datasets, the strict data policy and restricted age range present a major drawback for using it as a normative sample for training the BrainAGE models. We therefore combined images from IXI, OASIS-3, Cam-CAN, SALD, and Enhanced NKI-RS databases to create an independent normative dataset of healthy adults. An overview of the datasets used in this study can be seen in Table 1.

**Table 1.** Characteristics of the datasets used in this study.

| Dataset | | N (sessions) | Age range | M ± SD | Males/females |
| --- | --- | --- | --- | --- | --- |
| UKB | | 36840 | 45–82 | 64.09±7.53 | 17557/19283 |
| IXI | | 547 | 19–86 | 48.12±16.61 | 242/305 |
| OASIS-3 | | 549 | 42–97 | 69.53±8.86 | 208/341 |
| Cam-CAN | | 652 | 18–88 | 54.30±18.59 | 322/330 |
| SALD | | 494 | 19–80 | 45.18±17.44 | 187/307 |
| NKIe | | 629 | 18–85 | 38.74±21.27 | 233/396 |
| ADNI | HC | 231 | 60–90 | 76.01±5.01 | 119/112 |
| | sMCI | 100 | 58–88 | 75.40±7.27 | 66/34 |
| | pMCI | 164 | 55–88 | 74.57±7.03 | 97/67 |
| | AD | 200 | 55–91 | 75.64±7.72 | 103/97 |
| SZ | HC | 108 | 20–59 | 32.16±9.99 | 68/40 |
| | positive | 33 | 19–65 | 35.54±11.18 | 19/14 |
| | negative | 31 | 18–54 | 35.54±11.17 | 15/16 |
| | disorg. | 23 | 19–59 | 35.39±11.13 | 14/9 |
| Synthetic dataset | | 20 (20) | 59–78 | 70.65±5.39 | 10/10 |

Additionally, 695 T1w images from Alzheimer’s Disease Neuroimaging Initiative (ADNI, https://adni.loni.usc.edu/) were used to validate the model on a sample including healthy control subjects (HC, *n* = 231, *M*_Age_ = 76.01±5.01), subjects with stable MCI (sMCI, *n* = 100, *M*_Age_ = 75.40±7.27), subjects who progressed from MCI to Alzheimer’s dementia (pMCI, *n* = 164, *M*_Age_ = 74.57±7.03), and AD patients (*n* = 200, *M*_Age_ = 75.64±7.72).

We also used a synthetic global neocortical atrophy dataset by Rusak et al. (2022), which was derived from ADNI baseline scans of 20 individuals without AD (*n* = 20; *M*_Age_ = 70.65±5.39) and used to simulate neocortical thinning, progressing from 0 to 0.1 mm or 1 mm thickness loss, with steps of 0.01 mm or 0.1 mm between the time points, respectively, resulting in total of 400 synthetic images.

In order to validate the (regional) model on clinical data, a sample of schizophrenia patients and healthy controls previously described in Nenadić et al. (2015) was used.

### Data acquisition

The T1-weighted MR images of the UKB dataset were acquired on Siemens Skyra 3 T scanners with a 32-channel head coil. A sagittal MPRAGE sequence was used with 1 mm isotropic resolution, inversion/repetition/echo time (TI/TR/TE) = 880/2000/na ms, flip angle (FA) = na, field-of-view (FOV) = 208×256×256, acceleration factor (R) = 2, acquisition time: 5 minutes (https://biobank.ctsu.ox.ac.uk/crystal/crystal/docs/brain_mri.pdf).

In the ADNI sample used, T1w images were acquired on 1.5 T scanners manufactured by Siemens, Philips, and GE with single-channel coils. The sagittal MPRAGE sequences typically had about 0.95 mm in-plane resolution and 1.2 mm slice-thickness, TI/TR/TE = 853-1000/2300-3000/na ms, FA = 8-9°, FOV = 240-260×240 mm, R = 1, acquisition time: 7:11– 9:38 minutes, (Jack et al., 2008; see also https://adni.loni.usc.edu/methods/mri-tool/mri-acquisition/).

The IXI sample comprises data from 3 different scanners (GE 1.5 T, Philips 1.5 T and 3 T) with about 0.95 mm in-plane resolution and 1.2 mm slice-thickness, TI/TR/TE time = na/9.600-9.813/4.603 ms, FA = 8°, FOV = 256 mm, R = na (https://brain-development.org/ixi-dataset).

The MRI data from the OASIS-3 dataset used in this study were acquired on two Siemens TIM Trio and one BioGraph mMR PET-MR 3T scanners with 20-channel head coil. The MPRAGE scans were acquired with (i) 1 mm isotropic TI/TR/TE = 1000/2400/3.16 ms, FA = 8°, FOV phase = 100%, R = 2 and (ii) 1.20×1.05×1.05 mm resolution, TI/TR/TE = 900/2300/2.95 ms, FA = 9°, FOV phase = 93.75 mm, R = 2 (LaMontagne et al., 2019).

In the SALD dataset, a sagittal MPRAGE sequence on a 3 T Siemens Trio MRI scanner was used with 1 mm isotropic resolution, TI/TR/TE = 900/1900/2.52 ms, FA = 9°, FOV = 256 mm, R = 2 (Wei et al., 2018).

The Cam-CAN dataset was obtained on a 3 T Siemens TIM Trio scanner with a 32-channel head coil, using an MPRAGE sequence with 1 mm isotropic resolution, TI/TR/TE = 900/2250/2.98 ms, FA = 9°, FOV = 256×240×192 mm, R = 2, acquisition time = 4:32 minutes (Taylor et al., 2017).

T1 MPRAGE images from the NKI Enhanced dataset were acquired on a 3 T Siemens TIM Trio scanner with 32-channel head coil with 1 mm isotropic resolution, TI/TR/TE = 1200/2500/3.5 ms, FA = 8° (https://fcon_1000.projects.nitrc.org/indi/pro/eNKI_RS_TRT/FrontPage.html).

We used a sample of schizophrenia patients and healthy controls previously described in Nenadić et al. (2015). The sagittal MPRAGE images were acquired on a 1.5 T Philips Gyroscan ASCII scanner with 1 mm isotropic resolution, TI/TR/TE = na/13/5 ms, FOV = 256 mm.

### Data preprocessing

The UKB T1-weighted images were segmented into grey matter (GM) and white matter (WM) and (affinely/non-linearly) normalised by using default preprocessing of the CAT12.7 Toolbox (Gaser et al., 2022) as standalone version compiled under Matlab 2019b and SPM12 (Wellcome Centre for Human Neuroimaging, https://www.fil.ion.ucl.ac.uk/spm/) massively parallelized on the JURECA High Performance Computing system (JSC). The preprocessing utilises the unified segmentation (Ashburner & Friston, 2005) to remove B0-inhomogeneities and create an initial segmentation that is used for (local) intensity scaling and adaptive non-local means denoising (Manjon et al., 2010). An adaptive maximum a posteriori (AMAP; Rajapakse et al., 1997) segmentation with a hidden Markov random field (Cuadra et al., 2005) and partial volume effect model (Tohka et al., 2004) is utilised to create the final segmentation. For the optional non-linear registration, the Shooting method (Ashburner & Friston, 2011) with modulation was used.

The synthetic dataset, IXI, Cam-CAN, NKIe, OASIS-3, ADNI, and the SZ sample were processed with the CAT12.8 version with the same parameters.

### Global BrainAGE models estimation

In order to keep the process as parsimonious as possible, a linear affine registration of tissue segments to the MNI152Nlin2009cAsym template was used as default, except for a single non-linearly registered GM test case. Eight combinations of single tissue class models (GM/WM) with varying parameterisations of spatial resolution (4 mm / 8 mm) and Gaussian smoothing (FWHM: 4 mm / 8 mm) were trained and processed in the same way. With the exception of the first testing condition (see the test conditions below), principal component analysis (PCA) using singular value decomposition was applied to all the models to regularise the data. We employed a GPR model instead of the previously used RVR (see Figure 1A for the comparison of performance of both algorithms). This GPR uses a linear covariance function, a constant mean function and a Gaussian likelihood function with the hyperparameters set to 100 for the constant mean function and –1 for the likelihood function (Rasmussen & Williams, 2006). A conjugate gradient method was used for numerical optimization in the GPR.

**Figure 1.**
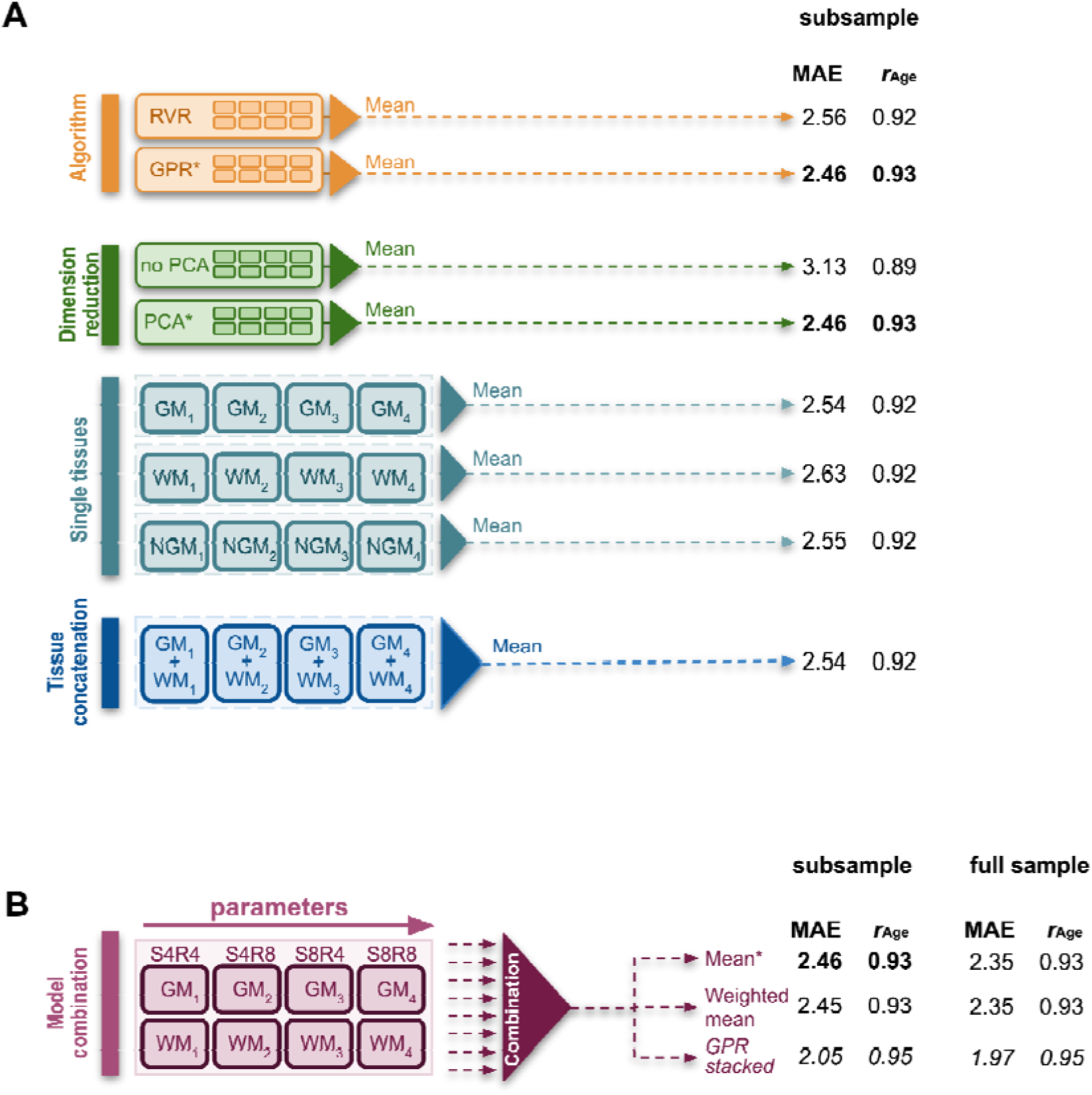
**A)** Various single models differing in spatial resolution (R: 4 mm / 8 mm), smoothing kernel (S: 4 mm / 8 mm), and tissue segments (GM, WM, NGM: non-linearly registered modulated GM) were estimated and combined. The results of model estimations combined by averaging are presented as age-bias corrected MAEs and the Pearson’s correlation coefficients between predicted- and chronological age for analyses run on a UKB 1 subsample. The comparison of the ML algorithms (RVR/GPR), as well as the effects of dimension reduction and the choice of brain tissues or their concatenation are shown. The results in bold represent the same combined model. **B)** Eight single models were ensembled by averaging, weighted averaging, or GPR stacking. The results are presented for the subsample and the full UKB sample.

A 10-fold cross-validation (CV) was performed with stratified chronological age to preserve the age distribution in each fold. The estimated brain age was age-bias adjusted by a linear term as described in Smith et al. (2019). Single tissue class models were assembled by averaging (unless stated otherwise).

The estimations were run in Matlab 2021a (Mathworks Inc., Natick, MA, USA).

### Regional BrainAGE models estimation

Regional BrainAGE models were trained on a large sample of healthy adult participants from 5 databases (IXI, OASIS-3, NKIe, SALD, Cam-CAN) and tested on the ADNI sample, spanning from healthy control subjects to AD patients, as well as on the SZ dataset. The training samples (AD: n = 1947, SZ: n = 2639) were age-matched to the ADNI and SZ test samples.

To obtain regional BrainAGE estimations, the GM segments were divided into 10 lobes (5 per hemisphere) derived from the Brain lobe atlas (Toro et al., 2009). Only GM segments were used as both previously mentioned diseases were shown to predominantly affect GM (Haijma et al., 2013; Wang et al., 2015). For each lobe, independent GPR models were estimated as described above. Only lobular regions were defined to support a useful amount of GM voxels for the models. A linear age-bias correction was used globally as described in Smith et al. (2019).

## Results

With respect to the change in the algorithm, we tested the novel machine learning framework performance under different conditions outlined in the following section, jointly reporting methods and results. We show the MAEs of the average-ensembled age-bias corrected models and Pearson’s correlation coefficients of the predicted age and the chronological age. Please see the Supplementary material (<u>Tables S1 & S2</u>) for the performance of all tested models without age-bias correction.

### I. The effect of principal component analysis

In this step, we tested the effect of implementing PCA using singular value decomposition, which was applied to the training data only with the resulting transformation used on the test sample to avoid data leakage. The performance of the model greatly improved with the drop from MAE of 3.13 years (*r*_Age_ = 0.89) to MAE of 2.46 years (*r*_Age_ = 0.93). All subsequent estimations on the full UKB dataset were therefore run on regularised data.

### II. Tissue class combination by model ensembling

Single tissue class models using only one linearly registered brain tissue class (either GM, WM, or non-linearly registered modulated GM), with varying parameterisations of spatial resolution (4 mm / 8 mm) and Gaussian smoothing (FWHM: 4 mm / 8 mm) were ensembled in order to minimise the MAE of the brain age prediction (Figure 1B). Three different ensembling approaches were tested, namely (i) their average performance, (ii) the weighted mean, and (iii) GPR stacking of the estimated models. The weighted mean was calculated with weighting w.r.t. squared MAE. The models with higher MAE were assigned a lower weight. A nested 10×10 CV was used in the GPR stacking, which utilised the predictions from the first level models as features for the second GPR learner.

We additionally tested the performance of mean-ensembled single-tissue-class models (either GM-, non-linearly registered modulated GM-, WM-, and concatenated GM- and WM vectors), which performed satisfactorily.

The results of ensembling different brain tissue classes are presented in Figure 1A and 1B for the subsample and the full sample, respectively. Model ensembling provides a more accurate BrainAGE estimation, with the most striking improvement obtained by stacking. Using the full UKB sample, GPR stacking resulted in 2.18 years of MAE for uncorrected BrainAGE prediction, MAE = 1.97 for linear age-bias correction as in Smith et al. (2019), and 2.34 years age-corrected as in Cole et al. (2018). Nevertheless, a simple mean-ensembled GM model was finally chosen for the application to the clinical datasets, as previous studies have shown that the models with highest performance accuracies are not the best in differentiating between (clinical) phenotypes (Bashyam et al., 2020).

### III. Sample dependence and the effect of training sample size

We additionally compared the performance of the algorithm between different subsamples of the UKB dataset. The subsamples were of comparable sizes and age/sex distributions and showed comparable results (see Table 2). We further tested the effect of varying training sample size by using 20% to 100% of the full dataset that showed only minor improvements in the performance of the algorithm (Supplementary Figure S1).

**Table 2.** Sample characteristics of different UKB subsamples with MAEs of the mean-ensembled model performance and Pearson’s correlation coefficients between the predicted and chronological age.

| Sample | N<sub>training</sub> | N<sub>testing</sub> | Males/Females | Age range | M ± SD | MAE [years] | $r_{\text{Age}}$ |
| --- | --- | --- | --- | --- | --- | --- | --- |
| UKB 1 | 6517 | 724 | 3503/3738 | 45–82 | 64.08±7.54 | **2.46** | **0.93** |
| UKB 2 | 6745 | 749 | 3618/3876 | 46–82 | 64.10±7.50 | 2.47 | 0.92 |
| UKB 3 | 6564 | 729 | 3406/3887 | 45–81 | 64.15±7.53 | 2.51 | 0.92 |
| UKB 4 | 6665 | 740 | 3523/3882 | 44–81 | 64.05±7.58 | 2.48 | 0.92 |
| UKB 5 | 6667 | 740 | 3507/3900 | 45–81 | **64.05±7.52** | 2.50 | 0.92 |
| UKB full | 33158 | 3682 | 17557/19283 | 44–82 | 64.09±7.53 | 2.35 | 0.93 |

### IV. Cross-scanner prediction reliability

We investigated the cross-site prediction by training the algorithm on a sample of 6631 T1-weighted MRI scans (using 90% of the sample to keep the size of training sample constant) from scanning site 1 (Cheadle, *M*_Age_ = 63.45±7.44, age range: 45–81) and tested its performance on a dataset from scanning site 3 (Newcastle, *M*_Age_ = 64.86±7.45, age range: 47–81), and vice versa. When predicting from scanning site 1 to 3, we obtained a MAE of 2.4 years (*r*_Age_ = 0.93), while the reverse resulted in a MAE of 2.48 years (*r*_Age_ = 0.92).

### Validation of the global model on external datasets

Due to the limited age range of the UKB sample, we trained the GM models on a previously described normative dataset, matching the age range of the training and validation samples. As we expect the most prominent differences in the GM tissue, only the results of the mean-ensembled GM model are presented in the following section.

#### I. Synthetic dataset of global neocortical thinning

We applied the global GM model to a synthetic dataset of global neocortical thinning from Rusak et al. (2022). The authors simulated neocortical thickness loss from 0 to 0.1 mm in steps of 0.01 mm per time point. The step reflects a yearly rate of cortical loss after the age of 70 (Fox & Schott, 2004; Fjell et al., 2009). They further simulated the hypothetical loss from 0.1 to 1 mm in steps of 0.1 mm which would altogether represent approximately 100 years of global neocortical thinning.

Our aim was to examine how the BrainAGE algorithm performs on simulated data with well-defined neocortical atrophy. We wanted to test whether the BrainAGE algorithm utilises the global neocortical thinning pattern for age prediction, and see to what extent the BrainAGE scores reflect the assumed yearly changes in neocortical thickness (Figure 2).

**Figure 2.**
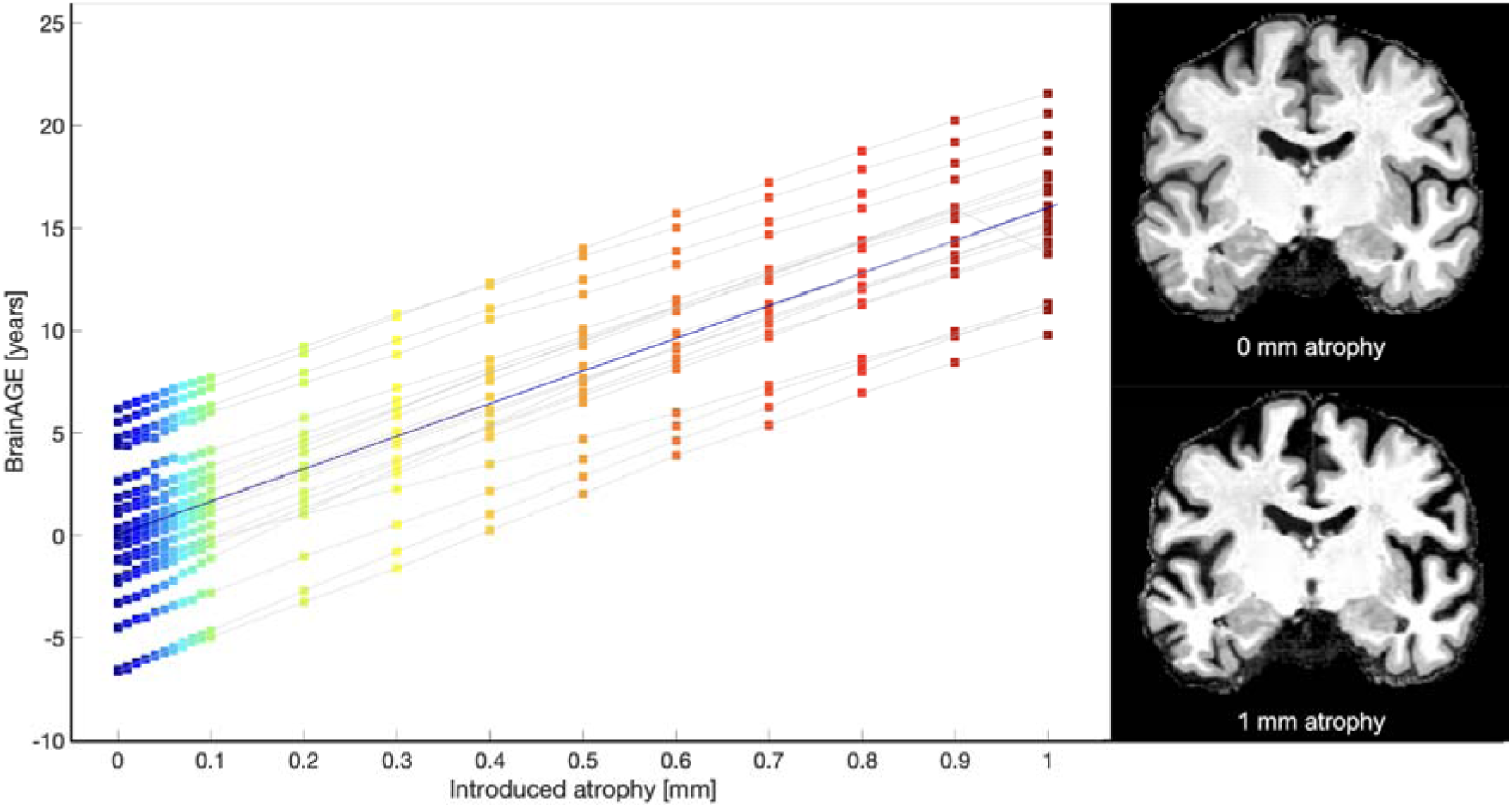
BrainAGE scores of the simulated neocortical thinning dataset with 0 and 1 mm atrophy examples on the right side.

As can be seen from the figure, BrainAGE scores reflect the loss of global cortical thickness, although the values do not represent the expected yearly rate of cortical thinning as only neocortical atrophy was simulated in the dataset.

#### II. Application to the ADNI sample

Furthermore, the global GM model was used to predict the BrainAGE in the ADNI sample of healthy controls, subjects with stable MCI and the ones who later progressed from MCI to AD, as well as AD patients. The ADNI test samples were matched with regard to age distribution of the normative database. The results are presented in Figure 3A.

**Figure 3.**
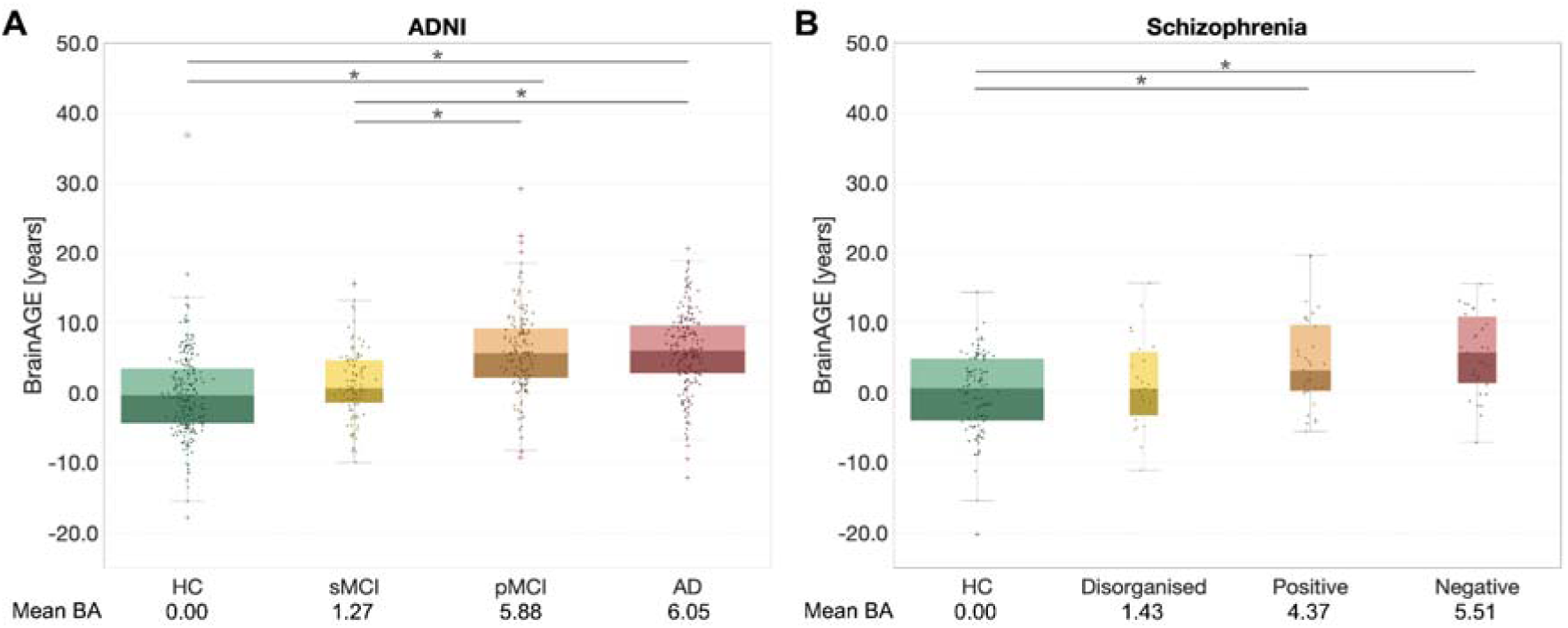
**A)** Mean BrainAGE scores for healthy individuals and patients with differing levels of neurocognitive impairment from the ADNI sample. **B)** Mean BrainAGE scores for healthy control participants and groups of patients with varying symptoms from the SZ sample.

One-way ANOVA showed statistically significant differences between the mean BrainAGE scores of the four groups (*F*(3, 691) = 55.69, *p* <.001). *Post hoc* comparisons using Tukey HSD test showed significant differences in mean BrainAGE scores of healthy controls and subjects with AD (*p*<.001), healthy controls and pMCI (*p*<.001), as well as sMCI and pMCI (*p*<.001) and sMCI and AD patients (*p*<.001) at baseline.

#### III. Application to the schizophrenia dataset

We applied the global GM model trained on a normative sample to the SZ dataset (Figure 3B). Statistically significant differences were found between the groups of patients with different symptoms (*F*(3, 191) = 9.64, *p* <.001). Tukey’s HSD *post hoc test* revealed statistically significant differences in mean BrainAGE scores of healthy controls and patients with negative (*p*<.001) and positive symptoms (*p* =.001), but not with the disorganised group (*p* =.715).

### Regional BrainAGE in health and disease

In order to support spatial specificity of the BrainAGE model, we developed a regional approach, parcellating the brain into 10 regions (5 per hemisphere). Separate GPR models were applied to a large sample of UKB and two clinical samples of AD and schizophrenia. The regional results from the large UKB sample can be seen in Table 3. Regional BrainAGE scores show an anterior-posterior pattern with higher frontal and lower occipital values, and a slight left-right difference with higher BrainAGE scores on the left compared to the right side. See <u>Tables S4 and S5</u> for the results without age-bias correction and with correction as in Cole (2020). The results of the models applied to AD and SZ patients can be seen in Figure 4.

**Figure 4.**
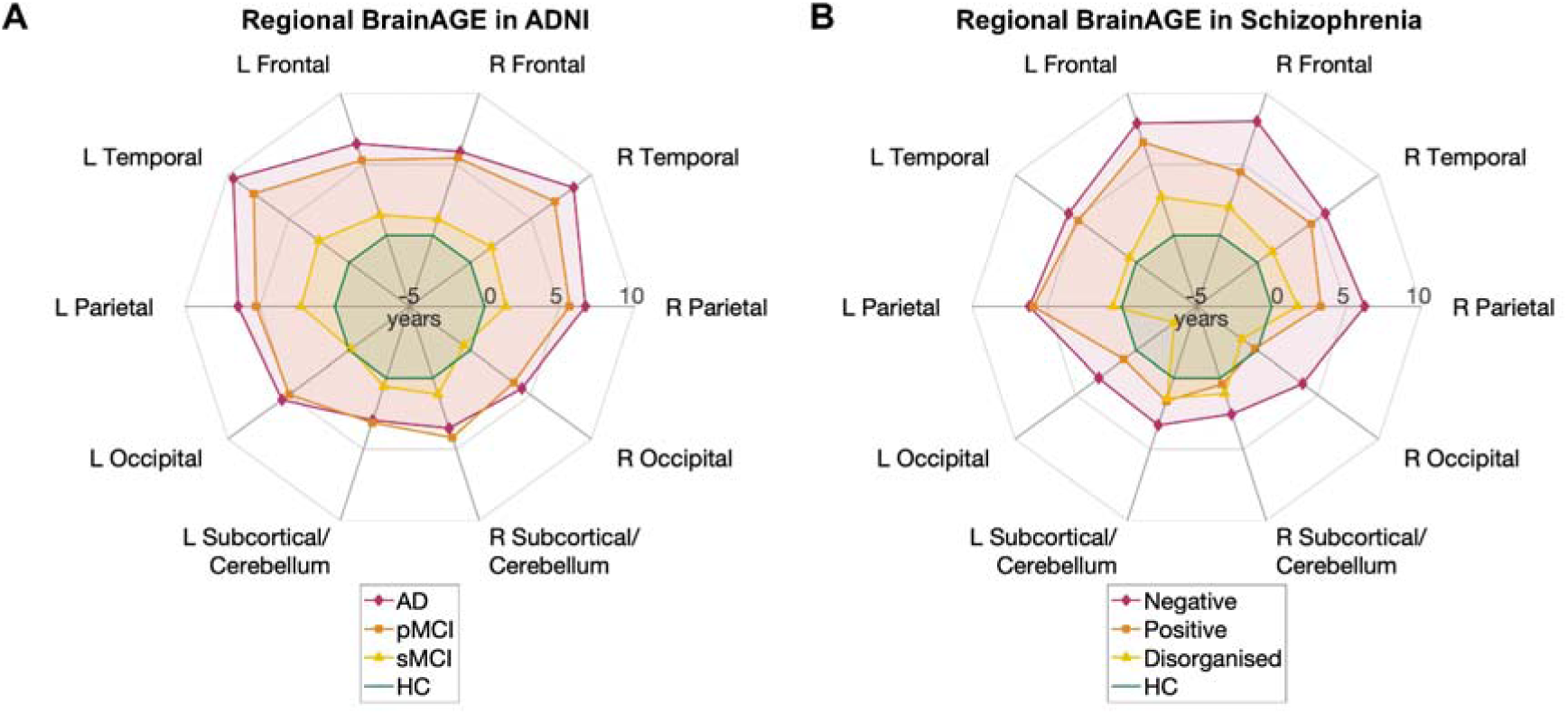
**A**) Regional BrainAGE scores in healthy control subjects (HC), patients with stable MCI (sMCI), progressive MCI (pMCI), and AD patients from the ADNI dataset. **B**) Regional BrainAGE scores for control subjects and three subgroups of SZ patients.

**Table 3.**
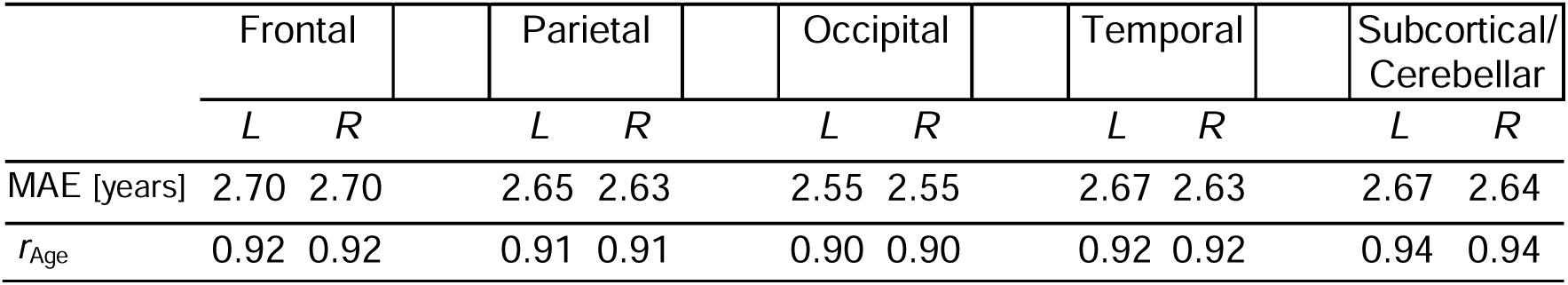
MAEs of the mean-ensembled model performance and Pearson’s correlation coefficients between the predicted and chronological age for regional BrainAGE models on the large sample of UKB with the linear age-bias correction as in Smith et al. (2019).

|  | Frontal |  | Parietal |  | Occipital |  | Temporal |  | Subcortical/<br>Cerebellar |  |
| --- | --- | --- | --- | --- | --- | --- | --- | --- | --- | --- |
|  | L | R | L | R | L | R | L | R | L | R |
| MAE [years] | 2.70 | 2.70 | 2.65 | 2.63 | 2.55 | 2.55 | 2.67 | 2.63 | 2.67 | 2.64 |
| $r_{Age}$ | 0.92 | 0.92 | 0.91 | 0.91 | 0.90 | 0.90 | 0.92 | 0.92 | 0.94 | 0.94 |

Higher BrainAGE scores can be observed especially in the temporal regions for AD and progressive MCI patients, whereas the sMCI group showed only minor differences overall and none in the occipital regions. The results in SZ patients are evident in the frontal, temporal, and parietal regions, most notably for the subjects with negative symptoms. The nearly unaffected disorganised group have only slightly increased brain age values mostly in frontal regions.

## Discussion

In our present work, we revised the BrainAGE model that was originally introduced more than a decade ago (Franke et al., 2010). The substitution of RVR with a GPR algorithm proved to be more stable in dealing with a large dataset, as demonstrated by extensive evaluation of the performance of the new framework under different conditions utilising the large neuroimaging sample of the UKB. We explored the contribution of grey and white matter tissue segments and their ensembles to BrainAGE prediction accuracy. By employing the model on the ADNI and SZ samples, we tested its external validity in dealing with pathological brain tissue changes, while the application of the model on a synthetic dataset showed how the algorithm reflects uniform decreases in global neocortical thickness. In addition, we developed a regional BrainAGE model to increase the spatial specificity of the BrainAGE prediction. We discuss the advantages and limitations of our new BrainAGE frameworks in the following sections.

### Dimension reduction & Training sample size

The results showed that one of the most important ways of improving the brain age gap estimation was the reduction of highly individual features that can be seen as “anatomical noise”. In our case this was conducted by smoothing, downsampling, and finally a PCA. As already noted in Franke et al. (2010), not all voxels of structural images are of equal importance to the brain age estimation, therefore, dimensionality reduction is one of the strategies to obtain more accurate results in case of highly dimensional feature space (Baecker et al., 2021; More et al., 2023). It does not only reduce the computational costs, but also prevent method-specific over-fitting of machine learning models (Franke & Gaser, 2019).

While RVR shows its superiority over GPR on smaller sample sizes (Baecker et al., 2021), training of a GPR model on a large enough sample yielded slightly better results than the RVR approach. However, increasing training sample size over 13,000 subjects resulted in plateauing of BrainAGE accuracies.

### Tissue class combination and model ensembling

Based on our analyses, a combination of models trained on affine registered GM and WM segments with varying preprocessing parameterization jointly contribute to the best prediction in our performance test on the UKB. Interestingly, the use of non-linearly registered GM data – typically used in VBM studies – did not result in a better performance in comparison to the models that utilised affine registered GM. We speculate that linear registration provides more spatial information of the individual brain anatomy and its atrophy pattern, being more analogous to raw brain anatomy used for deep learning methods, whereas the segmentation (with its denoising and bias-correction steps) helps to remove protocol-based image properties (Ashburner & Friston, 2000).

An important factor that strongly improved the prediction accuracy of BrainAGE estimation was combining various single BrainAGE models. Model ensembling is an effective way of improving the brain age estimation as it serves as a regularisation step to prevent overfitting and thus improves prediction accuracy (van Veen et al., 2015). Even though we ensembled models that were overall quite similar, we believe this is not a strong drawback since even similar models still provide additional variation (van Veen et al., 2015) and our results show improvement without significantly increased computational costs. As can be seen from our results, employing GPR stacking substantially decreased the MAE.

Additional improvements of age prediction may be possible by ensembling more independent brain age models, for example by using different image modalities (e.g. DWI, T2-weighted MR, rs-fMRI), morphometric metrics (e.g. thickness, sulcal depth), or other brain-age frameworks (e.g. Cole et al., 2018; Smith et al., 2019). Although (slight) improvements by adding modalities were shown before (e.g. Cole, 2020; Niu et al., 2020), we focused only on broadly available T1-weighted imaging data and a standard VBM paradigm in order to keep our framework parsimonious. Moreover, the inclusion of multiple modalities typically increases the accuracy of prediction, but does not lead to greater discrimination of phenotypes of interest (Jirsaraie et al., 2023). Nevertheless, the inclusion of other modalities may be useful where structural changes are not as prominent.

### Generalisability and validity

Our results show a comparable performance of the algorithm based on training and testing on different UKB subsamples. A repeated 10 x 10 CV performance demonstrated high consistency between the predicted BrainAGE scores across different folds and repetitions (<u>Table S3</u> in the supplementary material). Moreover, the cross-scanner prediction within two UKB sample sites shows consistent performance. Further testing of generalisability of the BrainAGE approach was done using the ADNI and SZ samples. The BrainAGE model showed meaningful differences between the diagnostic groups.

The application of the algorithm to the synthetic global neocortical atrophy dataset proved that the BrainAGE algorithm interprets cortical thinning as ageing, however, the obtained values do not reflect the expected yearly changes. This discrepancy is probably due to the properties of the synthetic dataset which reflects only the neocortical tissue loss, but does not simulate changes in white matter, ventricles, cerebellum, deep grey matter, hippocampus and amygdala (Rusak et al., 2022), which are structures that also drive brain age estimation (Levakov et al., 2020). Despite the artificial character of simulated atrophy pattern, such datasets allow for testing specific structural changes (Aubert-Broche et al., 2006, Lerch et al., 2005, Rusak et al., 2022) and their interaction with the brain age algorithms. Simulated datasets can potentially serve as a complementary approach to explainable AI by providing information on the influence of anatomical characteristics on brain age prediction.

Overall, the revised BrainAGE algorithm provides promising and generalisable results. Future studies might benefit from harmonising data over multiple imaging studies (Pomponio et al., 2020), thus allowing the application of brain age prediction to more diverse data.

### Regional BrainAGE

Global BrainAGE models fail to grasp neurodegenerative- and psychiatric disease-specific structural brain changes (Gianchandani et al., 2023; Kaufmann et al., 2019). In order to improve spatial specificity of the BrainAGE model, we developed a regional BrainAGE approach, estimating brain age in 5 lobular brain regions per hemisphere. The regional predictions were on average not as accurate as the global measure, which is expected given the lower information provided by a single lobe. Nevertheless, the regional BrainAGE scores provided meaningful information about brain ageing on clinical samples. The application to the ADNI sample showed an increase of BrainAGE scores in the temporal lobe of AD patients and MCI subjects converting to AD, corresponding to the implication of (medial) temporal lobe in AD (Migliaccio & Cacciamani, 2022). The results from the SZ sample were in accordance with the findings of the previous study on the same sample, where Nenadić et al. (2015) found the most prominent neocortical thinning in the negative symptoms group and the least in the disorganised group. Similarly, to the studies of Kaufmann et al. (2019) and Zhu et al. (2023), SZ patients in our sample had the highest BrainAGE scores in the frontal and temporal parts. Additional studies are needed to further validate the regional BrainAGE model in various clinical groups.

## Conclusion

Our BrainAGE algorithm underwent a thorough revision that enabled its application on the very large UK Biobank dataset. Moreover, we extended the framework to support regional brain age estimation and increase the spatial specificity of the method. Application of the BrainAGE models on subjects with varying levels of neuropsychiatric- and neurocognitive impairment showcases the model’s clinical validity and the BrainAGE scores from a synthetic dataset reflect a meaningful pattern according to known neocortical thickness loss with age. Future research can profit from the use of synthetic data to better understand the inner workings of ML algorithms.

## Acknowledgements

This research has been conducted using data from UK Biobank under the application number 41655.

Data were provided in part by OASIS-3: Longitudinal Multimodal Neuroimaging: Principal Investigators: T. Benzinger, D. Marcus, J. Morris; NIH P30 AG066444, P50 AG00561, P30 NS09857781, P01 AG026276, P01 AG003991, R01 AG043434, UL1 TR000448, R01 EB009352. AV-45 doses were provided by Avid Radiopharmaceuticals, a wholly owned subsidiary of Eli Lilly.

The work was supported by Carl Zeiss Stiftung as a part of the IMPULS project (IMPULS P2019-01-006), the Federal Ministry of Science and Education (BMBF) under the frame of ERA PerMed (Pattern-Cog ERAPERMED2021-127) and the Marie Skłodowska-Curie Innovative Training Network (SmartAge 859890 H2020-MSCA-ITN2019).

The authors gratefully acknowledge the computing time granted by the JARA Vergabegremium and provided on the JARA Partition part of the supercomputer JURECA at Forschungszentrum Jülich.

## Authorship contribution statement

**Polona Kalc**: Writing – original draft. Visualisation. **Robert Dahnke**: Writing – review & editing, Visualisation, Data curation, Software. **Felix Hoffstaedter**: Data curation, Writing – review & editing. **Christian Gaser**: Conceptualization, Methodology, Formal analysis, Software, Writing – review & editing, Visualisation, Supervision, Funding acquisition.

## Data availability statement

The data used for this work was obtained from the UK Biobank Resource (Project Number 41655). Due to the nature of the data sharing agreement, we are not allowed to publish the data.

The ADNI dataset is publicly available at https://ida.loni.usc.edu/login.jsp?project=ADNI and IXI at https://brain-development.org/ixi-dataset/, Cam-CAN is available under https://camcan-archive.mrc-cbu.cam.ac.uk/dataaccess/, OASIS-3 can be found on https://www.oasis-brains.org/#data, Enhanced NKI-RS dataset is available at https://fcon_1000.projects.nitrc.org/indi/pro/eNKI_RS_TRT/FrontPage.html, and the SALD dataset can be downloaded from https://fcon_1000.projects.nitrc.org/indi/retro/sald.html. The synthetic dataset from Rusak et al. (2022) is available at the CSIRO data access portal.

## Supplementary material

**Table S1.** Mean absolute errors (and Pearson’s corr. coefficients between predicted and chronological age) for different age-bias corrections of the BrainAGE estimated on the UKB 1 subsample.

|  | *No age-bias*<br>*correction* | *Linear age-bias*<br>*correction*<br>*(Smith et al., 2019)* | *Age-bias*<br>*correction*<br>*(Cole, 2020)* |
| --- | --- | --- | --- |
| *Mean* | 3.03<br>(0.86) | 2.46<br>(0.93) | 3.48<br>(0.86) |
| *Weighted mean* | 3.02<br>(0.87) | 2.45<br>(0.93) | 3.47<br>(0.87) |
| *GPR stacking* | 2.26<br>(0.93) | 2.05<br>(0.95) | 2.44<br>(0.93) |

**Table S2.** Mean absolute errors (and Pearson’s corr. coefficients between predicted and chronological age) for different age-bias corrections of the BrainAGE estimated on the full UKB sample.

|  | *No age-bias*<br>*correction* | *Linear age-bias*<br>*correction*<br>*(Smith et al., 2019)* | *Age-bias*<br>*correction*<br>*(Cole, 2020)* |
| --- | --- | --- | --- |
| *Mean* | 2.80<br>(0.89) | 2.35<br>(0.93) | 3.14<br>(0.89) |
| *Weighted mean* | 2.80<br>(0.89) | 2.35<br>(0.93) | 3.13<br>(0.89) |
| *GPR stacking* | 2.19<br>(0.93) | 1.97<br>(0.95) | 2.34<br>(0.93) |

**Table S3.**
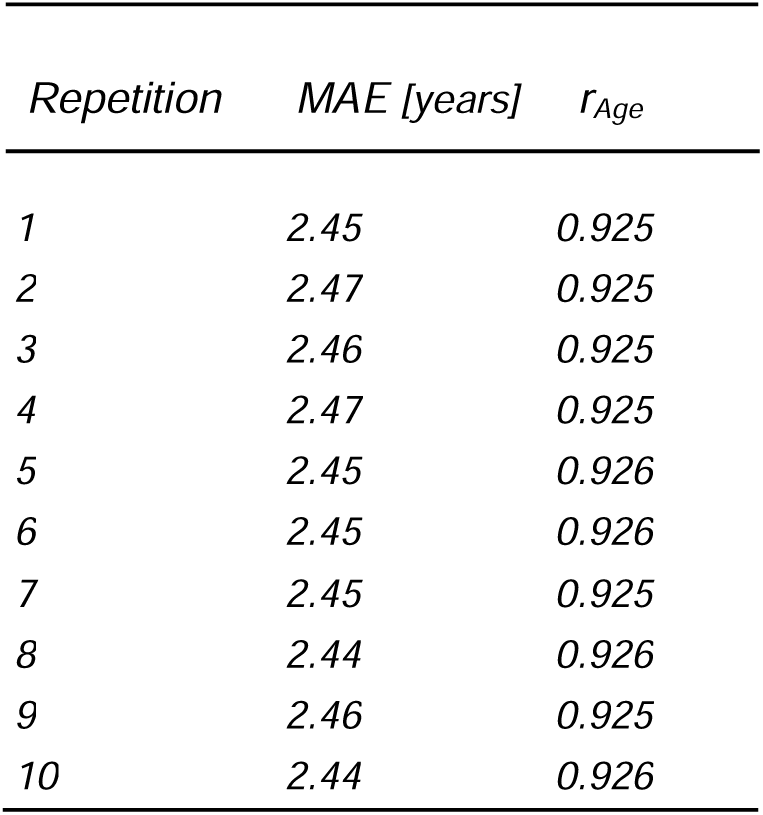
Mean absolute errors and Pearson’s correlation coefficient between brain age estimation and chronological age averaged across all CV folds for 10 repetitions (in the repeated 10 x 10-fold CV).

**Table S4.**
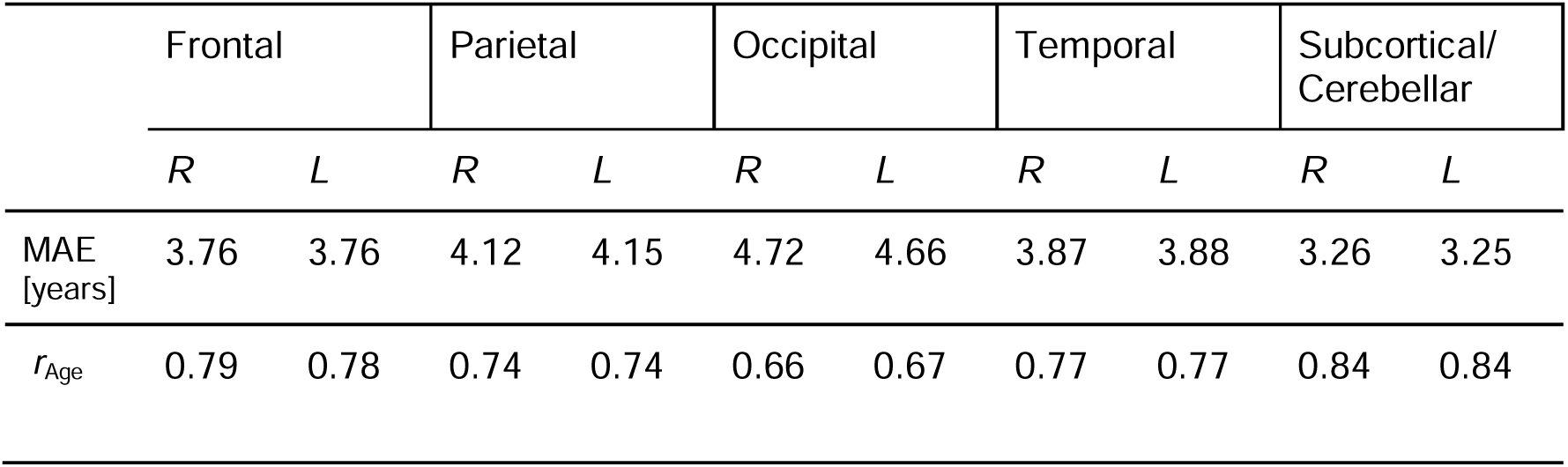
MAEs of the mean-ensembled model performance on the large UKB sample and Pearson’s correlation coefficients between the predicted and chronological age for regional BrainAGE models **without age-bias correction**.

|  | Frontal |  | Parietal |  | Occipital |  | Temporal |  | Subcortical/<br>Cerebellar |  |
| --- | --- | --- | --- | --- | --- | --- | --- | --- | --- | --- |
|  | R | L | R | L | R | L | R | L | R | L |
| MAE [years] | 3.76 | 3.76 | 4.12 | 4.15 | 4.72 | 4.66 | 3.87 | 3.88 | 3.26 | 3.25 |
| $r_{Age}$ | 0.79 | 0.78 | 0.74 | 0.74 | 0.66 | 0.67 | 0.77 | 0.77 | 0.84 | 0.84 |

**Table S5.** *MAEs of the mean-ensembled model performance on the large UKB sample and Pearson’s correlation coefficients between the predicted and chronological age for regional BrainAGE models with age-bias correction* as in Cole (2020).

| Frontal | Parietal | Occipital | Temporal | Subcortical/<br>Cerebellar |
| --- | --- | --- | --- | --- |

|  | *R* | *L* | *R* | *L* | *R* | *L* | *R* | *L* | *R* | *L* |
| --- | --- | --- | --- | --- | --- | --- | --- | --- | --- | --- |
| MAE [years] | 5.21 | 5.22 | 5.08 | 5.12 | 4.92 | 4.94 | 5.08 | 5.15 | 5.10 | 5.16 |
| $r_{\text{Age}}$ | 0.79 | 0.78 | 0.74 | 0.74 | 0.66 | 0.67 | 0.77 | 0.77 | 0.84 | 0.84 |

**Figure S1.**
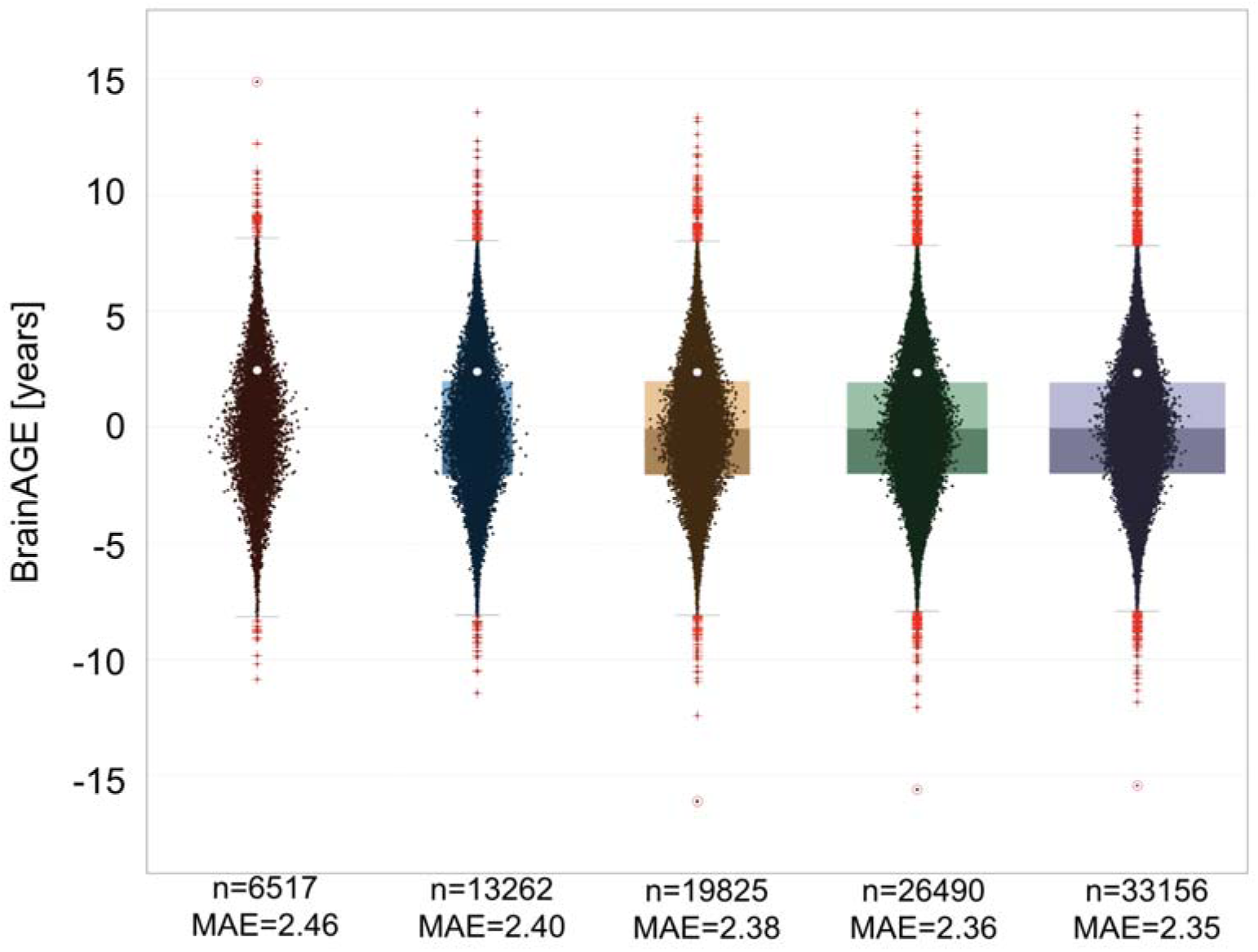
Mean absolute errors for different training sample sizes from the UKB sample.

## References

Abrol, A., Fu, Z., Salman, M., Silva, R., Du, Y., Plis, S., & Calhoun, V. (2021). Deep learning encodes robust discriminative neuroimaging representations to outperform standard machine learning. Nature Communications, 12(1), 353. 10.1038/s41467-020-20655-6

Ashburner, J., & Friston, K. J. (2000). Voxel-based morphometry--the methods. NeuroImage, 11, 805–821. doi:10.1006/nimg.2000.0582

Ashburner, J., & Friston, K. J. (2005). Unified segmentation. NeuroImage, 26(3), 839–851. 10.1016/j.neuroimage.2005.02.018

Ashburner, J., & Friston, K. J. (2011). Diffeomorphic registration using geodesic shooting and Gauss–Newton optimisation. NeuroImage, 55(3), 954–967. 10.1016/j.neuroimage.2010.12.049

Aubert-Broche, B., Evans, A. C., & Collins, D. L. (2006). A new improved version of the realistic digital brain phantom. NeuroImage 32, 138–145. 10.1016/j.neuroimage.2006.03.052

Baecker, L., Dafflon, J., Costa, P. F., GarciaLDias, R., Vieira, S., Scarpazza, C., Calhoun, V. D., Sato, J. R., Mechelli, A., & Pinaya, W. H. L. (2021). Brain age prediction: A comparison between machine learning models using regionL and voxelLbased morphometric data. Human Brain Mapping, 42(8), 2332–2346. 10.1002/hbm.25368

Bashyam, V. M., Erus, G., Doshi, J., Habes, M., Nasrallah, I. M., Truelove-Hill, M., Srinivasan, D., Mamourian, L., Pomponio, R., Fan, Y., Launer, L. J., Masters, C. L., Maruff, P., Zhuo, C., Völzke, H., Johnson, S. C., Fripp, J., Koutsouleris, N., Satterthwaite, T. D., … Davatzikos, C. (2020). MRI signatures of brain age and disease over the lifespan based on a deep brain network and 14 468 individuals worldwide. Brain, 143(7), 2312–2324. 10.1093/brain/awaa160

Beheshti, I., Mishra, S., Sone, D., Khanna, P., & Matsuda, H. (2020). T1-weighted MRI-driven Brain Age Estimation in Alzheimer’s Disease and Parkinson’s Disease. Aging and Disease, 11(3), 618. 10.14336/AD.2019.0617

Beheshti, I., Ganaie, M. A., Paliwal, V., Rastogi, A., Razzak, I., & Tanveer, M. (2022). Predicting Brain Age Using Machine Learning Algorithms: A Comprehensive Evaluation. IEEE Journal of Biomedical and Health Informatics, 26(4), 1432–1440. 10.1109/JBHI.2021.3083187.

Bethlehem, R. A. I., Seidlitz, J., White, S. R., Vogel, J. W., Anderson, K. M., Adamson, C., Adler, S., Alexopoulos, G. S., Anagnostou, E., Areces-Gonzalez, A., Astle, D. E., Auyeung, B., Ayub, M., Bae, J., Ball, G., Baron-Cohen, S., Beare, R., Bedford, S. A., Benegal, V., … Alexander-Bloch, A. F. (2022). Brain charts for the human lifespan. Nature, 604(7906), 525–533. 10.1038/s41586-022-04554-y

Chen, T., & Ren, J. (2009). Bagging for Gaussian process regression. Neurocomputing, 72(7–9), 1605–1610. 10.1016/j.neucom.2008.09.002

Cole, J. H. (2020). Multimodality neuroimaging brain-age in UK Biobank: Relationship to biomedical, lifestyle, and cognitive factors. Neurobiology of Aging, 92, 34–42. 10.1016/j.neurobiolaging.2020.03.014

Cole, J. H., & Franke, K. (2017). Predicting Age Using Neuroimaging: Innovative Brain Ageing Biomarkers. Trends in Neurosciences, 40(12), 681–690. 10.1016/j.tins.2017.10.001

Cole, J. H., Leech, R., Sharp, D. J., for the Alzheimer’s Disease Neuroimaging Initiative. (2015). Prediction of brain age suggests accelerated atrophy after traumatic brain injury. Annals of Neurology, 77(4), 571–581. 10.1002/ana.24367

Cole, J. H., Ritchie, S. J., Bastin, M. E., Valdés Hernández, M. C., Muñoz Maniega, S., Royle, N., Corley, J., Pattie, A., Harris, S. E., Zhang, Q., Wray, N. R., Redmond, P., Marioni, R. E., Starr, J. M., Cox, S. R., Wardlaw, J. M., Sharp, D. J., & Deary, I. J. (2018). Brain age predicts mortality. Molecular Psychiatry, 23(5), 1385–1392. 10.1038/mp.2017.62

Cuadra, M. B., Cammoun, L., Butz, T., Cuisenaire, O., & Thiran, J.-P. (2005). Comparison and validation of tissue modelization and statistical classification methods in T1-weighted MR brain images. IEEE Transactions on Medical Imaging, 24(12), 1548–1565. 10.1109/TMI.2005.857652

Eickhoff, C. R., Hoffstaedter, F., Caspers, J., Reetz, K., Mathys, C., Dogan, I., Amunts, K., Schnitzler, A., Eickhoff, S. B. (2021). Advanced brain ageing in Parkinson’s disease is related to disease duration and individual impairment. Brain Communications, 3(3), 1–12. 10.1093/braincomms/fcab191

Fjell, A. M., Walhovd, K. B., Fennema-Notestine, C., McEvoy, L. K., Hagler, D. J., Holland, D., Brewer, J. B., Dale, A. M. (2009). One-year brain atrophy evident in healthy aging. J Neurosci, 29(48), 15223–31. doi: 10.1523/JNEUROSCI.3252-09.2009.

Fox, N. C., & Schott, J. M. (2004). Imaging cerebral atrophy: normal ageing to Alzheimer’s disease. Lancet, 363, 392–394. doi: 10.1016/S0140-6736(04)15441-X

Franke, K., Clarke, G. D., Dahnke, R., Gaser, C., Kuo, A. H., Li, C., Schwab, M., & Nathanielsz, P. W. (2017). Premature Brain Aging in Baboons Resulting from Moderate Fetal Undernutrition. Frontiers in Aging Neuroscience, 9. 10.3389/fnagi.2017.00092

Franke, K., & Gaser, C. (2012). Longitudinal Changes in Individual *BrainAGE* in Healthy Aging, Mild Cognitive Impairment, and Alzheimer’s Disease. GeroPsych, 25(4), 235–245. 10.1024/1662-9647/a000074

Franke, K., & Gaser, C. (2019). Ten Years of BrainAGE as a Neuroimaging Biomarker of Brain Aging: What Insights Have We Gained? Frontiers in Neurology, 10, 789. 10.3389/fneur.2019.00789

Franke, K., Gaser, C., Manor, B., & Novak, V. (2013). Advanced BrainAGE in older adults with type 2 diabetes mellitus. Frontiers in Aging Neuroscience, 5. 10.3389/fnagi.2013.00090

Franke, K., Luders, E., May, A., Wilke, M., & Gaser, C. (2012). Brain maturation: Predicting individual BrainAGE in children and adolescents using structural MRI. NeuroImage, 63(3), 1305–1312. 10.1016/j.neuroimage.2012.08.001

Franke, K., Ziegler, G., Klöppel, S., & Gaser, C. (2010). Estimating the age of healthy subjects from T1-weighted MRI scans using kernel methods: Exploring the influence of various parameters. NeuroImage, 50(3), 883–892. 10.1016/j.neuroimage.2010.01.005

Gaser, C., Dahnke, R., Thompson, P. M., Kurth, F., & Luders, E. (2022). *CAT - A Computational Anatomy Toolbox for the Analysis of Structural MRI Data* [Preprint]. biorXiv. 10.1101/2022.06.11.495736

Gianchandani, N., Dibaji, M., Ospel, J., Vega, F., Bento, M., MacDonald, M. E., & Souza, R. (2023). A voxel-level approach to brain age prediction: A method to assess regional brain aging (arXiv:2310.11385). arXiv. http://arxiv.org/abs/2310.11385

Gong, W., Beckmann, C. F., Vedaldi, A., Smith, S. M., & Peng, H. (2020). *Optimising a Simple Fully Convolutional Network (SFCN) for accurate brain age prediction in the PAC 2019 challenge* [Preprint]. Neuroscience. 10.1101/2020.11.10.376970

Hahn, T., Fisch, L., Ernsting, J., Winter, N. R., Leenings, R., Sarink, K., Emden, D., Kircher, T., Berger, K., & Dannlowski, U. (2021). From ‘loose fitting’ to high-performance, uncertainty-aware brain-age modelling. Brain, 144(3), e31–e31. 10.1093/brain/awaa454

Haijma, S. V., Van Haren, N., Cahn, W., Koolschijn, P. C. M. P., Hulshoff Pol, H. E., & Kahn, R. S. (2013). Brain Volumes in Schizophrenia: A Meta-Analysis in Over 18 000 Subjects. Schizophrenia Bulletin, 39(5), 1129–1138. 10.1093/schbul/sbs118

Jack, C. R., Bernstein, M. A., Fox, N. C., Thompson, P., Alexander, G., Harvey, D., Borowski, B., Britson, P. J., L. Whitwell, J., Ward, C., Dale, A. M., Felmlee, J. P., Gunter, J. L., Hill, D. L. G., Killiany, R., Schuff, N., Fox-Bosetti, S., Lin, C., Studholme, C., … ADNI Study. (2008). The Alzheimer’s disease neuroimaging initiative (ADNI): MRI methods. Journal of Magnetic Resonance Imaging, 27(4), 685–691. 10.1002/jmri.21049

Jirsaraie, R. J., Gorelik, A. J., Gatavins, M. M., Engemann, D. A., Bogdan, R., Barch, D. M., & Sotiras, A. (2023). A systematic review of multimodal brain age studies: Uncovering a divergence between model accuracy and utility. Patterns, 4(4), 100712. 10.1016/j.patter.2023.100712

Jülich Supercomputing Center. (2018). JURECA: Data Centric and Booster Modules implementing the Modular Supercomputing Architecture at Jülich Supercomputing Centre Journal of large-scale research facilities, 7, A182. 10.17815/jlsrf-7-182

Kaufmann, T., Van Der Meer, D., Doan, N. T., Schwarz, E., Lund, M. J., Agartz, I., Alnæs, D., Barch, D. M., Baur-Streubel, R., Bertolino, A., Bettella, F., Beyer, M. K., Bøen, E., Borgwardt, S., Brandt, C. L., Buitelaar, J., Celius, E. G., Cervenka, S., … Westlye, L. T. (2019). Common brain disorders are associated with heritable patterns of apparent aging of the brain. Nature Neuroscience, 22(10), 1617–1623. 10.1038/s41593-019-0471-7

LaMontagne, P. J., Benzinger, T. Ls., Morris, J. C., Keefe, S., Hornbeck, R., Xiong, C., Grant, E., Hassenstab, J., Moulder, K., Vlassenko, A. G., Raichle, M. E., Cruchaga, C., & Marcus, D. (2019). OASIS-3: Longitudinal Neuroimaging, Clinical, and Cognitive Dataset for Normal Aging and Alzheimer Disease [Preprint]. Radiology and Imaging. 10.1101/2019.12.13.19014902

Levakov, G., Rosenthal, G., Shelef, I., Raviv, T. R., & Avidan, G. (2020). From a deep learning model back to the brain—Identifying regional predictors and their relation to aging. Hum. Brain Mapp., 41(12), 3235–3252. doi: 10.1002/hbm.25011

Lombardi, A., Monaco, A., Donvito, G., Amoroso, N., Bellotti, R., & Tangaro, S. (2021). Brain Age Prediction With Morphological Features Using Deep Neural Networks: Results From Predictive Analytic Competition 2019. Frontiers in Psychiatry, 11, 619629. 10.3389/fpsyt.2020.619629

Lerch, J. P. & Evans, A. C. (2005). Cortical thickness analysis examined through power analysis and a population simulation. NeuroImage, 24, 163–173. 10.1016/j.neuroimage.2004.07.045

Luders, E., Cherbuin, N., & Gaser, C. (2016). Estimating brain age using high-resolution pattern recognition: Younger brains in long-term meditation practitioners. NeuroImage, 134, 508–513. 10.1016/j.neuroimage.2016.04.007

Manjon, J., Coupe, P., Marti-Bonmati, L., Collins, D.L., & Robles, M. (2010). Adaptive Non-Local Means Denoising of MR Images With Spatially Varying Noise Levels. Journal of Magnetic Resonance Imaging, 31, 192–203. 10.1002/jmri.22003

Migliaccio, R., & Cacciamani, F. (2022). The temporal lobe in typical and atypical Alzheimer disease. In Handbook of Clinical Neurology (Vol. 187, pp. 449–466). Elsevier. 10.1016/B978-0-12-823493-8.00004-3

More, S., Antonopoulos, G., Hoffstaedter, F., Caspers, J., Eickhoff, S. B., & Patil, K. R. (2023). Brain-age prediction: A systematic comparison of machine learning workflows. NeuroImage, 270, 119947. 10.1016/j.neuroimage.2023.119947

Nenadić, I., Yotter, R., Sauer, H., & Gaser, C. (2015). Patterns of cortical thinning in different subgroups of schizophrenia. The British Journal of Psychiatry, 206(6), 479–483. doi:10.1192/bjp.bp.114.148510

Nenadić, I., Dietzek, M., Langbein, K., Sauer, H., & Gaser, C. (2017). BrainAGE score indicates accelerated brain aging in schizophrenia, but not bipolar disorder. Psychiatry Research: Neuroimaging, 266, 86–89. 10.1016/j.pscychresns.2017.05.006

Niu, X., Zhang, F., Kounios, J., & Liang, H. (2020). Improved prediction of brain age using multimodal neuroimaging data. Human Brain Mapping, 41(6), 1626–1643. 10.1002/hbm.24899

Peng, H., Gong, W., Beckmann, C. F., Vedaldi, A., & Smith, S. M. (2021). Accurate brain age prediction with lightweight deep neural networks. Medical Image Analysis, 68, 101871. 10.1016/j.media.2020.101871

Pomponio, R., Erus, G., Habes, M., Doshi, J., Srinivasan, D., Mamourian, E., Bashyam, V., Nasrallah, I. M., Satterthwaite, T. D., Fan, Y., Launer, L. J., Masters, C. L., Maruff, P., Zhuo, C., Völzke, H., Johnson, S. C., Fripp, J., Koutsouleris, N., Wolf, D. H., … Davatzikos, C. (2020). Harmonization of large MRI datasets for the analysis of brain imaging patterns throughout the lifespan. NeuroImage, 208, 116450. 10.1016/j.neuroimage.2019.116450

Rajapakse, J. C., Giedd, J. N., & Rapoport, J. L. (1997). Statistical approach to segmentation of single-channel cerebral MR images. IEEE Transactions on Medical Imaging, 16(2), 176–186. 10.1109/42.563663

Rasmussen, C. E., & Williams, C. K. I. (2006). Gaussian processes for machine learning. MIT Press.

Rogenmoser, L., Kernbach, J., Schlaug, G., & Gaser, C. (2018). Keeping brains young with making music. Brain Structure and Function, 223(1), 297–305. 10.1007/s00429-017-1491-2

Rusak, F., Santa Cruz, R., Lebrat, L., Hlinka, O., Fripp, J., Smith, E., Fookes, C., Bradley, A. P., & Bourgeat, P. (2022). Quantifiable brain atrophy synthesis for benchmarking of cortical thickness estimation methods. Medical Image Analysis, 82, 102576. 10.1016/j.media.2022.102576

Smith, S. M., Vidaurre, D., Alfaro-Almagro, F., Nichols, T. E., & Miller, K. L. (2019). Estimation of brain age delta from brain imaging. NeuroImage, 200, 528–539. 10.1016/j.neuroimage.2019.06.017

Taylor, J. R., Williams, N., Cusack, R., Auer, T., Shafto, M. A., Dixon, M., Tyler, L. K., Cam-CAN, & Henson, R. N. (2017). The Cambridge Centre for Ageing and Neuroscience (Cam-CAN) data repository: Structural and functional MRI, MEG, and cognitive data from a cross-sectional adult lifespan sample. NeuroImage, 144, 262–269. 10.1016/j.neuroimage.2015.09.018

Tohka, J., Zijdenbos, A., & Evans, A. (2004). Fast and robust parameter estimation for statistical partial volume models in brain MRI. NeuroImage, 23(1), 84–97. 10.1016/j.neuroimage.2004.05.007

Toro, R., Chupin, M., Garnero, L., Leonard, G., Perron, M., Pike, B., Pitiot, A., Richer, L., Veillette, S., Pausova, Z., & Paus, T. (2009). Brain volumes and Val66Met polymorphism of the BDNF gene: local or global effects?. Brain Struct Funct, 213, 501–509. 10.1007/s00429-009-0203-y

van Veen, H. J., The Dat, L. N., Segnini, A. (2015). Kaggle Ensembling Guide. https://mlwave.com/kaggle-ensembling-guide/

Varzandian, A., Razo, M. A. S., Sanders, M. R., Atmakuru, A., & Di Fatta, G. (2021). Classification-Biased Apparent Brain Age for the Prediction of Alzheimer’s Disease. Frontiers in Neuroscience, 15, 673120. 10.3389/fnins.2021.67312

Wang, W.-Y., Yu, J.-T., Liu, Y., Yin, R.-H., Wang, H.-F., Wang, J., Tan, L., Radua, J., & Tan, L. (2015). Voxel-based meta-analysis of grey matter changes in Alzheimer’s disease. Translational Neurodegeneration, 4(1), 6. 10.1186/s40035-015-0027-z

Wei, D., Zhuang, K., Ai, L., Chen, Q., Yang, W., Liu, W., Wang, K., Sun, J., & Qiu, J. (2018). Structural and functional brain scans from the cross-sectional Southwest University adult lifespan dataset. Scientific Data, 5(1), 180134. 10.1038/sdata.2018.134

Zhu, J.-D., Wu, Y.-F., Tsai, S.-J., Lin, C.-P., & Yang, A. C. (2023). Investigating brain aging trajectory deviations in different brain regions of individuals with schizophrenia using multimodal magnetic resonance imaging and brain-age prediction: A multicenter study. Translational Psychiatry, 13(1), 82. 10.1038/s41398-023-02379-5

